# Synaptic silencing affects the density and complexity of oligodendrocyte precursor cells in the adult mouse hippocampus

**DOI:** 10.1101/2020.09.23.309682

**Authors:** Irene Chacon-De-La-Rocha, Gemma L. Fryatt, Andrea D. Rivera, Laura Restani, Matteo Caleo, Olivier Raineteau, Diego Gomez-Nicola, Arthur M. Butt

## Abstract

Oligodendrocyte progenitor cells (OPCs) are responsible for generating oligodendrocytes, the myelinating cells of the CNS. Life-long myelination is promoted by neuronal activity and is essential for neural network plasticity and learning. OPCs are known to contact synapses and it is proposed that neuronal synaptic activity in turn regulates OPC proliferation and differentiation. To examine this in the adult, we performed unilateral injection of the synaptic blocker botulinum neurotoxin A (BoNT/A) into the hippocampus of adult mice. We confirm BoNT/A cleaves SNAP-25 in the CA1 are of the hippocampus, which has been proven to block neurotransmission. Notably, synaptic silencing by BoNT/A significantly decreased OPC density and caused their shrinkage, as determined by immunolabelling for the OPC marker NG2. Inhibition of synaptic activity resulted in an overall decrease in the number of OPC processes, as well as a decrease in their lengths and branching frequency. These data indicate that synaptic activity is important for maintaining adult OPC numbers and cellular integrity, which is relevant to pathological scenarios characterized by decreased synaptic activity.

## Introduction

Oligodendrocyte precursor cells (OPCs) are a significant population of cells in the adult brain with the fundamental function of life-long generation of oligodendrocytes, which is required to myelinate new connections formed in response to new life experiences and to replace myelin lost through natural ‘wear and tear’ and disease^1–3^. OPCs are identified by their expression of the proteoglycan NG2 (Cspg4)^4,5^ and have a complex morphology, extending processes to contact neuronal synapses and nodes of Ranvier, sensing neuronal glumatergic and GABAergic activity^6–11^. Several lines of evidence indicate neuronal synaptic activity regulates OPC proliferation and differentiation^12–15^. Optogenetic studies have shown that neuronal activity stimulates OPC proliferation and differentiation in the cortex^16^. Furthermore, motor learning has been shown to drive OPC differentiation and myelination of newly formed neuronal connections, and failure to generate new oligodendrocytes impairs learning ability^17,18^. Hence, it is proposed that decreased synaptic activity may result in disruption of OPC regenerative capacity in neurodegenerative diseases, such as Multiple Sclerosis (MS) and Alzheimer’s disease (AD)^19,20^.

The synaptic protein SNAP-25 is necessary for synaptic vesicle fusion and its cleavage by botulinum toxins (BoNTs) blocks neurotransmitter release and synaptic signalling^21^. Developmental myelination is inhibited by ablation of neuronal SNAP-25 either by BoNT/A in cell culture^22^, or genetically in vivo^23^, but the effects of synaptic activity on adult OPCs had not been resolved. In the present study, we investigated the effects of prolonged synaptic silencing on adult OPCs, by local injection of botulinum toxin A (BoNT/A) into the mouse hippocampal CA1 region, which has been shown to produce a sustained blockade of synaptic transmission via cleavage of the synaptic protein SNAP-25^21,24,25^. Our results demonstrate that synaptic silencing by BoNT/A decreases the density and complexity of OPCs, leading to overall cell morphological atrophy. These results provide direct evidence about the role of synaptic activity in the maintenance of the population of OPCs, with relevance for understanding any pathological process leading to progressive demyelination.

## Results

### Synaptic silencing with BoNT/A causes a reduction in OPC numerical density and size in the CA1 region of the hippocampus

The long-term effects of synaptic silencing on adult OPCs have not been previously studied. We addressed this by injecting adult mice into the left hippocampus with BoNT/A (1 nM solution, 0.2 μl), which causes persistent blockade of hippocampal synaptic activity for up to 120 days^24,25^. The hippocampus was analysed 2 weeks post-injection, a time at which BoNT/A-mediated SNAP-25 cleavage was evident in the CA1 area of inoculated side (Fig. 1), confirming previous studies^24,25^.

**Figure 1.**
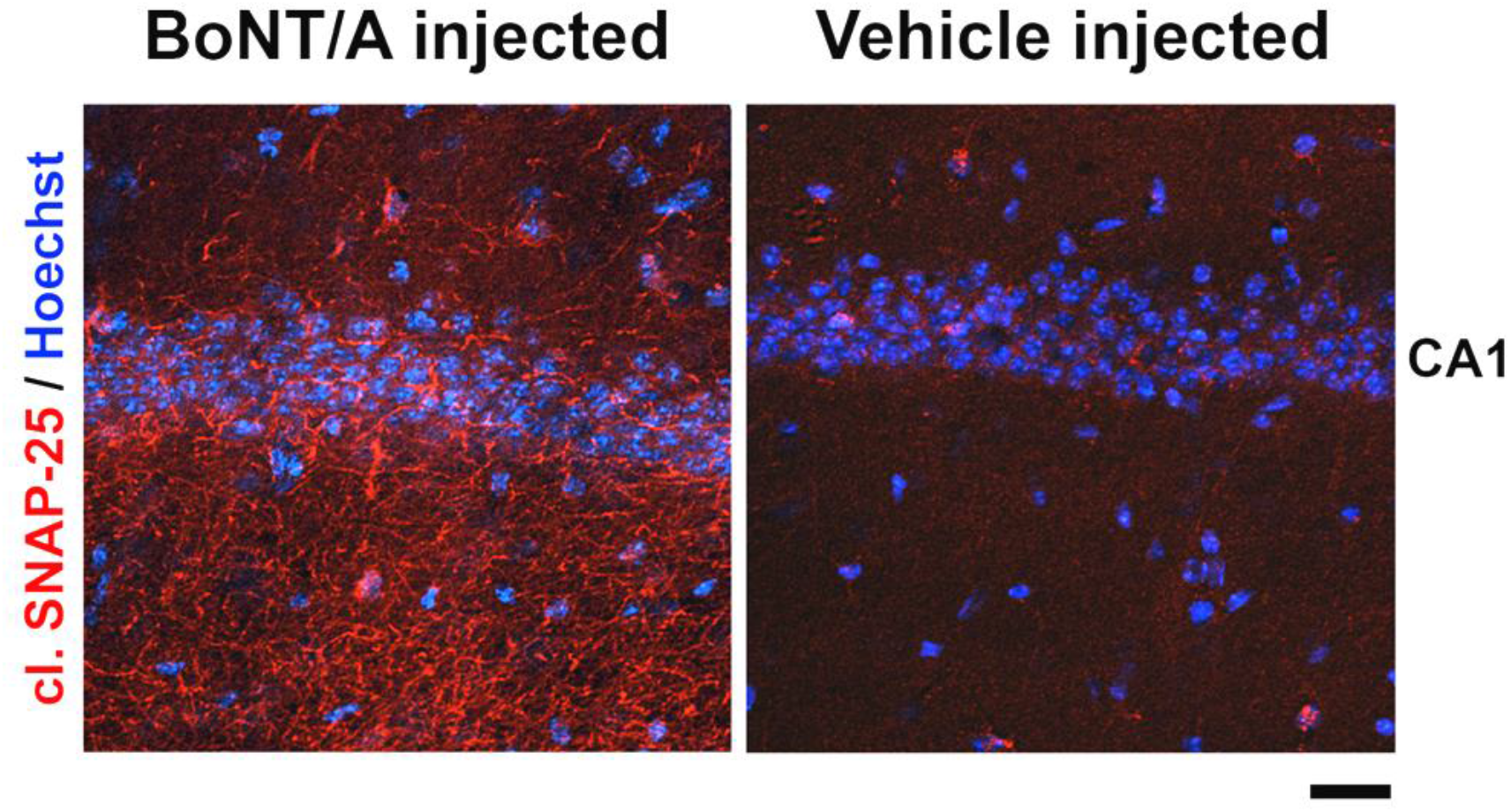
BoNT/A cleaves SNAP-25 in the hippocampus. Fluorescence micrographs of the CA1 are of the adult mouse hippocampus following stereotaxical injection of BoNT/A or vehicle. Sections are immunolabelled with an antibody specific for BoNT/A-truncated SNAP-25 (red), which has been thoroughly characterized in previous studies^24,25^, and counterstained with Hoechst nuclear dye. Scale bar = 100 μm.

Next, we used immunolabelling for NG2 to identify OPCs^6^, and we focused on the CA1 area as OPCs had been previously shown to form synapses with neurones and respond to synaptic signalling^7,9^ Confocal images demonstrate an evident decrease in NG2 immunostaining 14 days after BoNT/A injection (Fig. 2A), compared to controls injected with vehicle (Fig. 2B). Quantification confirmed OPC density was decreased following BoNT/A injection compared to controls (Fig. 2C; *p*<0.01, unpaired t-test). Moreover, mapping the process domains of individual OPCs (Fig. 2D, E) demonstrated these were significantly reduced following synaptic silencing (Fig. 2F; *p*<0.001, unpaired t-test). The data show that synaptic silencing results in a decline in the overall number of OPC and those that persist display a marked shrinkage.

**Figure 2.**
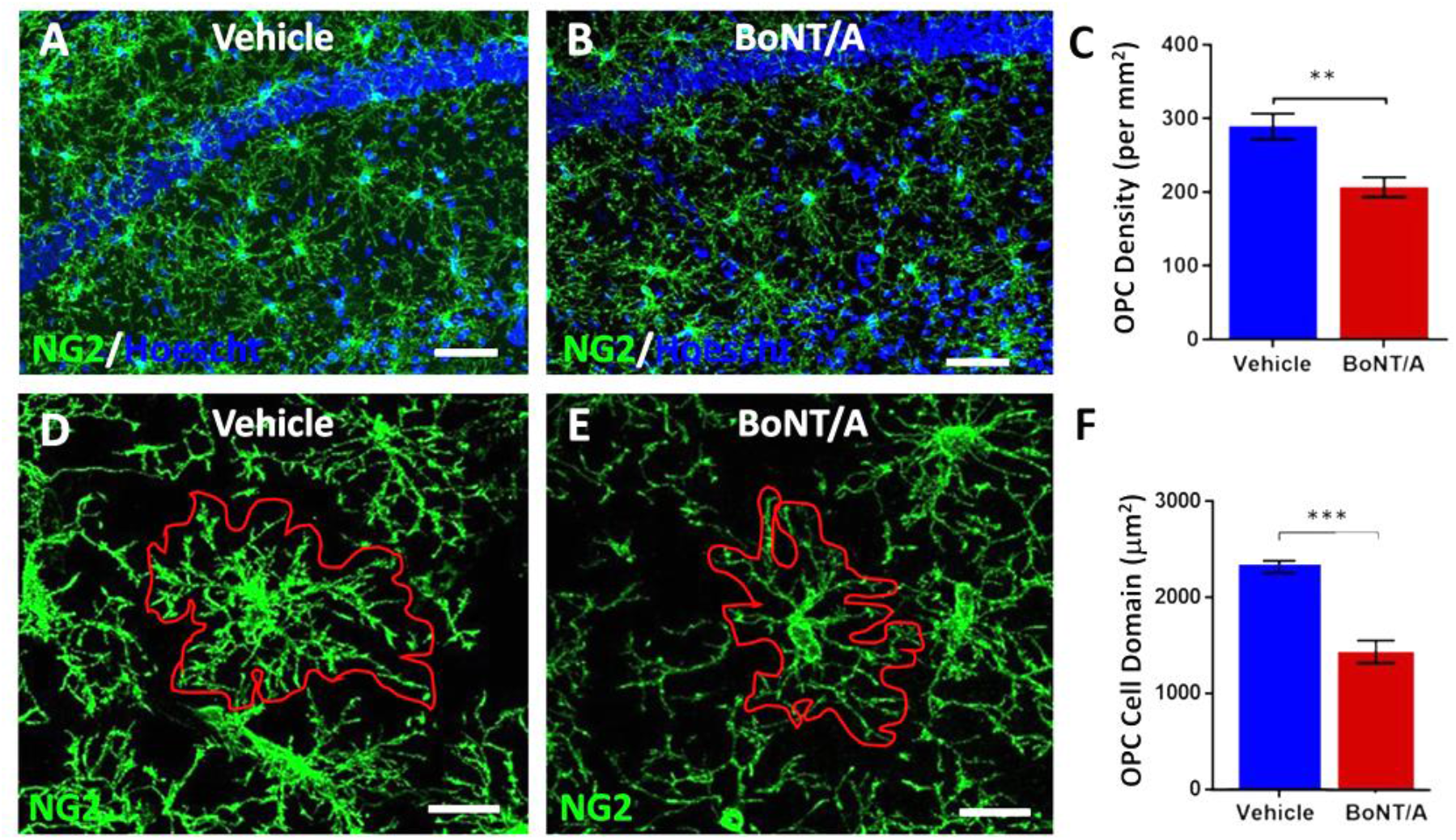
Effect of synaptic silencing on adult OPC. Confocal analysis of OPCs in hippocampus following injection with BoNT/A to silence synaptic activity, or vehicle in controls. (**A-C**) Immunolabelling for NG2 (green) and counterstained with Hoechst nuclear dye (blue), illustrating the overall distribution of OPCs in the CA1 area of the hippocampus following injection of vehicle (A) or BoNT/A (B), together with bar graph of the numerical density of OPCs in the CA1 area of the hippcampus (C). Scale bars = 50 μm. Data expressed as mean ± SEM (n=4 animals per group); \*\**p*<0.01, unpaired t-tests. (**D-F**) High magnification of individual NG2 immunolabelled OPCs in the CA1 area of the hippocampus illustrating their process domains (red lines) following injection of vehicle (D) or BoNT/A (E), together with bar graph of OPC cellular domains (F). Scale bars = 20 μm. Data expressed as mean ± SEM (n=4 animals per group); \*\*\**p*<0.001, unpaired t-test.

### BoNT/A decreases the morphological complexity of OPCs

Adult OPCs respond to pathology by changes in their morphology^26^, and in the adult hippocampus they react to toxic activation of glutamatergic receptors by extending a greater number of short process^5^. We therefore examined OPC morphology in detail using Neurolucida (Fig. 3) and Sholl analysis (Fig. 4). Overall, OPCs had a characteristic complex morphology in vehicle injected controls, with on average 12 primary processes that extended radially for 50-100 μm from a central cell body (Fig. 3A). In comparison, OPCs displayed overall atrophy following BoNT/A injection, with an evident decrease in cellular complexity (Fig. 3B); the shrinkage of the OPC process field following BoNT/A treatment is amplified in the y-plane (upper insets, Fig. 3A, B), and in the x-plane OPCs in controls are seen to have a dense network of branching processes, which is stunted following synaptic silencing (lower insets, Fig. 3A, B). Multiple parameters of process complexity were quantified (Fig. 3C-E; *n*=12 cells per group, Mann Whitney tests). The number of processes extending from the cell body was decreased following BoNT/A injection, ranging from 2-12, compared to 8-25 in controls, but these differences were not statistically significant (Fig. 3C). In contrast, there were significant decreases in the total length of processes from each cell (Fig. 3D), as well as the number of process terminals, or end points (Fig. 3E), and the number of branch points along processes (nodes) (Fig. 3F). The results indicate a primary effect of BoNT/A on cellular branching and this was examined further by Sholl analysis (Fig. 4A; *n*=12 cells for each group, two-way ANOVA followed by Sidak’s multiple comparisons test). The data indicate that following synaptic silencing the most marked changes in OPC processes were within 20-30 μm of the cell body, with significant decreases in the number of nodes, or branch points (Fig. 4B), as well as the number of process terminals (Fig. 3C), and the lengths of processes (Fig. 4D). In addition, we analysed the processes length in the different branch orders, identifying that synaptic silencing primarily caused shrinkage of the distal branches, and the maximum branch order was decreased to 13 compared to 15 in controls (Fig. 4E). The morphological analyses demonstrate that OPCs undergo significant morphological atrophy following synaptic silencing in the hippocampus.

**Figure 3.**
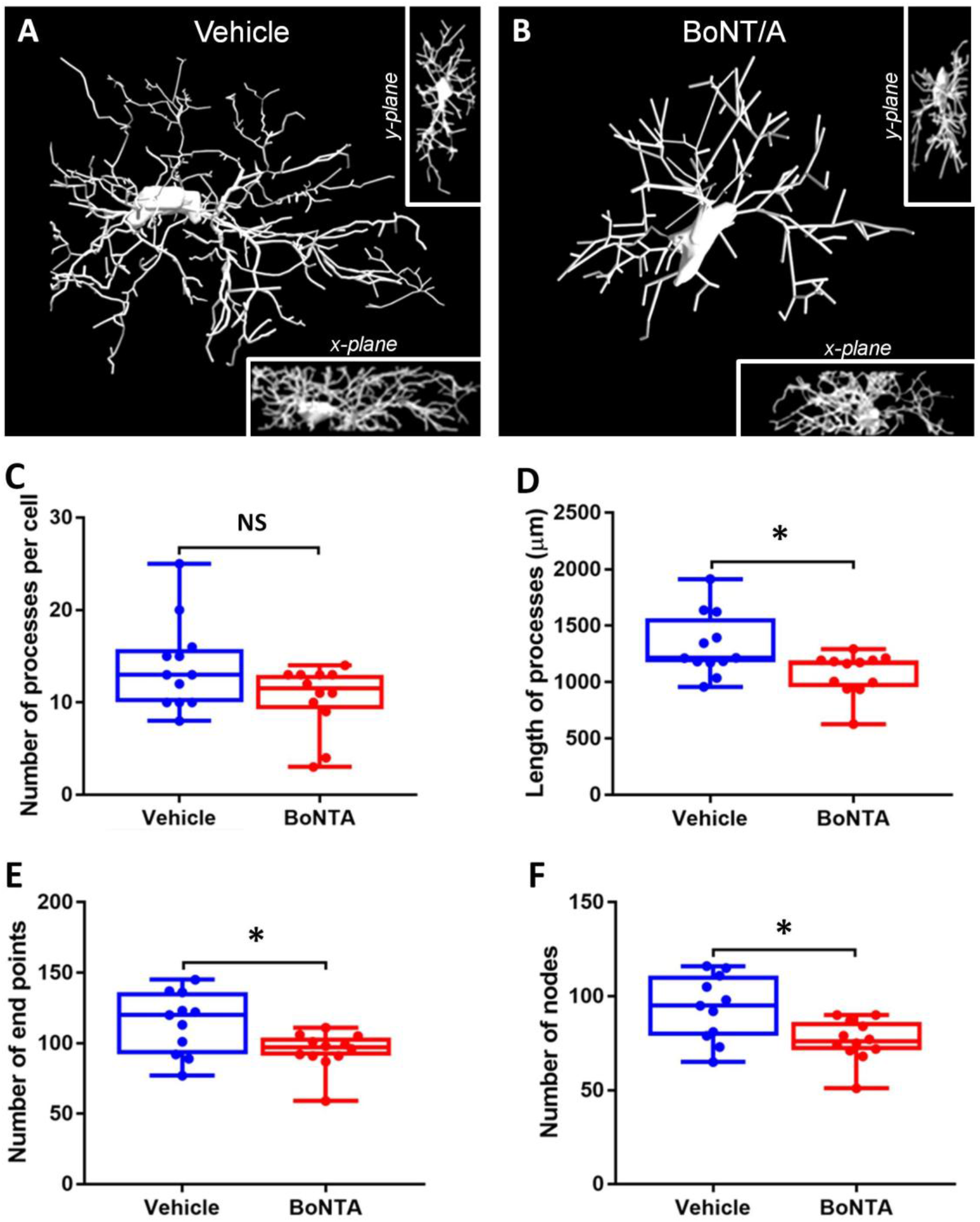
Effects of synaptic silencing on OPC morphology. Confocal images of hippocampus sections were immunolabelled for NG2 following injection with BoNT/A to silence synaptic activity, or vehicle in controls. (A, B) Confocal images were analysed using Neurolucida 360 (MBF Bioscience), to generate 3D images (main panels in A,B); insets illustrate the cell in the x- and y-planes. (C-F) Box-whisker plots of the Number of processes per cell (C), Process Lengths (D), Number of process terminals, or end points (E), and the Number of branch points, or nodes (F). Data are Mean ± SEM (*n*=11 or 12 cells from 4 sections from 4 animals per group; \**p*<0.05, Mann-Whitney test.

**Figure 4.**
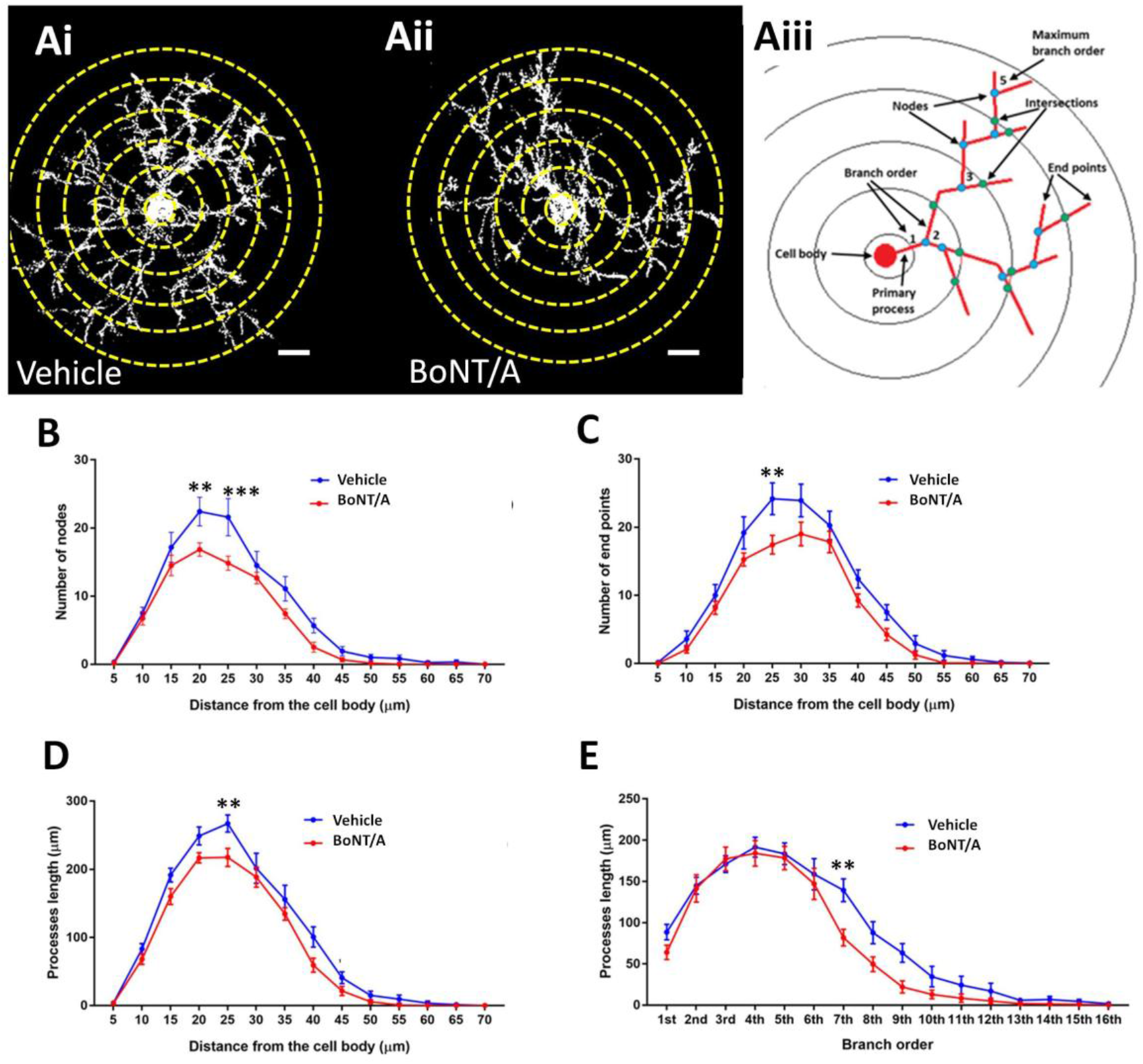
Sholl analysis of OPC process morhology following synaptic silencing. (A) Isosurface rendering to illustrate 3D morphology of OPCs (generated with Volocity software, PerkinElmer), obtained following NG2 immunolabelled cells in CA1 area of the hippocampus following injection of BoNT/A (Ai) or vehicle (Aii). The schematic representation in (Aiii) illustrates parameters measured, where the concentric circles (termed Sholl shells) are placed at 5 mm apart, with the cell body in the middle (yellow lines in Ai, Aii)l the points of process branching are termed nodes (blue dots), and the points where the processes intersect the Sholl shells are termed intersections (green dots) (adapted from Sholl 1953 and Rietveld et al. 2015). (B-E) Graphs of the Number of nodes or branch points (B), Number of process terminals or end points (C), Processes lengths (D), and Length of processes of different branch orders (E). Data are Mean ± SEM (*n* = 12 cells from 4 animals per group; \*\**p*<0.01, \*\*\**p*<0.001, two-way ANOVA followed by Sidak’s multiple comparisons test.

## Discussion

An important feature of OPCs is that they form synapses with neurones and sense neuronal synaptic activity via their neurotransmitter receptors^7,9^. Several lines of evidence support that synaptic signalling regulates OPC proliferation and differentiation^12–15^. Here, we show that injection of BoNT/A into the hippocampus, which cleaves SNAP-25 and silences synaptic signalling^24^, results in a decrease in OPC density and a decrease in OPC morphological complexity. Notably, similar changes in OPCs have been reported as an early event associated with synaptic dysregulation in AD-like pathology in mice^20^. The results support a role for synaptic signalling in maintaining adult OPC numbers and integrity, which is relevant to neuropathologies in which neuronal activity is altered, such as AD and MS.

BoNT/A is a member of the family of botulinum toxins (BoNTs A-G) that inhibit neurotransmission by blocking vesicular neurotransmitter release^21^. BoNTs enter presynaptic terminals mainly via activity-dependent synaptic endocytosis and cleave proteins of the SNARE complex, which is necessary for synaptic vesicles fusion^27^. It has been demonstrated that infusion of BoNT/A into the mouse hippocampus cleaves SNAP-25 and prevents neuronal synaptic activity in the CA1 area^24^. Glia and their precursors may express the genetic machinery for vesicular release, but at far lower levels than neurones^28^, and the balance of evidence indicates that glia express SNAP-23 and not SNAP-25^29–31^. *In vitro*, oligodendrocytes express SNAP-23 when they mature^31,32^. In the hippocampus, SNAP-25 is absent from glial processes and is concentrated in neuronal terminals^30^. In addition, BoNT/A cleaves SNAP-25, but not SNAP-23^33^, and BoNT/A has little direct effect on astrocytes^34^. Overall, the results of the present study are consistent with BoNT/A not affecting glia directly and specifically silencing neuronal synapses^24,25^, which induces marked changes in adult OPCs.

Synaptic silencing by BoNT/A resulted in a decrease in the overall density of OPCs, suggesting OPC self-renewal and/or survival was dependent on neuronal activity. The decrease in OPCs following synaptic silencing indicates their population was not fully replenished by increased proliferation, which is the usual response of OPCs to CNS insults^26,35^. This supports evidence that blockade of neuronal activity with TTX reduces OPC proliferation and survival^36^, whereas stimulation of neuronal activity stimulates OPC proliferation^16^. A key finding of our study is that synaptic silencing by BoNT/A has a marked effect on OPC morphology, most notably resulting in cell atrophy, marked by decreased branching and shrinkage of the distal processes. The resulting overall decrease in OPC coverage could be caused by a retraction of OPC processes from silent synapses, which is consistent with a study in cultured developing cerebellar slices showing blockade of neuronal activity and glutamatergic signalling reduces OPC process extension and branching^14^. In addition, time-lapse imaging demonstrates that NG2^+^ OPCs are highly dynamic, surveying their local environment with motile filopodia and responding rapidly to changes in their environment^37^. Notably, atrophy of OPC is an early sign of AD-like pathology and is concomitant with dysregulation of synaptic signaling in the 3xTg-AD mouse^20^. Indeed, changes in OPC morphology is a characteristic of neuropathology in human AD and mouse models of the disease^38–41^. Overall, our data demonstrate that blockade of synaptic signalling in the adult hippocampus results in OPC morphological atrophy and a decline in their numbers.

In summary, this study demonstrates for the first time that inhibition of synaptic neurotransmission by BoNT/A results in reduced numbers of OPCs and marked cellular shrinkage. We confirm that BoNT/A causes cleavage of SNAP-25, which has been shown unequivocally to preferentially suppress hippocampal excitatory activity^24,25^. GABAergic synapses are reported to be less sensitive to BoNT/A, due to SNAP-25 being expressed only at very low levels in GABAergic synapses^42^. Nonetheless, SNAP-25 expression has been reported in GABAergic neurones of the adult hippocampus and GABAergic transmission is disrupted in neurones cultured from SNAP-25 knock out mice^43,44^.Furthermore, OPCs in the hippocampus receive synaptic inputs from both glutamatergic and GABAergic neurones^7,9^, indicating that both neurotransmitter systems may influence the observed effects of BoNT/A on adult OPCs. However, we observed a decrease in OPC density following BoNT/A, whereas blockade of GABAergic signalling appears to increase OPC numbers in cortical slices^12,15^. In addition, we observed striking morphological changes in OPCs, which has been reported following blockade of glutamatergic signalling in cerebellar slice cultures^14^. In conclusion, the effects of BoNT/A on OPCs is consistent with a primary effect on inhibition of excitatory synaptic input onto OPCs. This study demonstrate that disruption of neuronal activity has a major negative impact on adult OPC numbers and integrity, which is relevant to multiple neuropathologies, including AD and MS.

## Materials and Methods

### Ethics statement

All procedures were performed in compliance with the EU Council Directive 2010/63/EU on the protection of animals used for scientific purposes, and approved by the Italian Ministry of Health (protocol # 346/2013-B). All surgical procedures were performed under deep anaesthesia and all efforts were made to ameliorate suffering of animals.

### Animals and Tissues

C57BL/6N mice were bred and group housed in the animal facility of the CNR Neuroscience Institute in Pisa (Italy). Mice aged 11-12 weeks were anesthetized by intraperitoneal injection of Avertin (2,2,2-tribromoethanol solution) (20 ml/kg) and a stereotaxically guided injection of BoNT/A (0.2 μl of a 1 nM solution) or vehicle (2% rat serum albumin in PBS) was made into the dorsal hippocampus using fine glass micropipettes, as previously described^24^. In all BoNT/A injected animals, recovery was uneventful and no overt behavioural abnormalities were observed. Two weeks after injections, mice were deeply anesthetized with chloral hydrate and perfused through the heart with freshly prepared 4% paraformaldehyde in 0.1 M phosphate buffer (PB), and the brains processed for immunohistochemistry (see below).

### Immunohistochemistry

Brains were post-fixed for 1 h at 4°C and brain sections (50 μm thick) were cut with a freezing microtome and stored at 4°C until use in cryoprotectant solution containing 25% sucrose and 3.5% glycerol in 0.05 M PB at pH 7.4. To detect SNAP-25 cleavage, we used an antibody specific for BoNT/A-truncated SNAP-25 which has been thoroughly characterized in previous studies^24,25^. Sections were blocked with 10% donkey serum in PBS containing 0.25% Triton X-100 and then incubated overnight at room temperature with the anti-cleaved SNAP-25 antibody (1:500 dilution). On the following day, sections were incubated with anti-rabbit Rhodamine Red X (1:500, Jackson ImmunoResearch), washed in PBS, counterstained with Hoechst dye and mounted with antifading agent. For NG2 immunostaining, sections were blocked with 0.5% bovine serum albumin (BSA) for 1-2 h, washed 3 times in PBS, and incubated overnight in the antibody solution which comprised of the primary antibody rabbit anti-NG2, 1:500 (Millipore), diluted in blocking solution containing 0.25% Triton-X. Tissues were then washed 3 times in PBS and incubated with appropriate fluorochrome secondary antibody diluted in blocking solution for 1-2h (AlexaFluor^®^ 488, AlexaFluor^®^ 568, 1:400, Life Technologies). Finally, sections were washed 3 times with PBS before being mounted on glass slides and covered with mounting medium and glass coverslips ready for imaging.

### Imaging and Analysis

Immunofluorescence images were captured using a Zeiss Axiovert LSM 710 VIS40S confocal microscope and maintaining the acquisition parameters constant to allow comparison between samples within the same experiment. Acquisition of images for cell counts was done with x20 objective. Images for OPC reconstruction were taken using x100 objective and capturing z-stacks formed by 80-100 single plains with an interval of 0.3 μm. OPC cell counts were performed in a constant field of view (FOV) of 200 μm x 200 μm and date expressed as OPC density per mm^2^. To determine the process domains (cell coverage) of OPCs, a line was drawn around the entire cell process field and the area within the line was measured using ImageJ and data were expressed as μm^2^. For morphological analysis, single OPCs were drawn in detail using Neurolucida 360 and their morphology was analysed using Neurolucida 360 explorer. For Sholl analysis, the interval between Sholl shells was 5μm; in addition, we analysed the processes length in the different branch orders (1^st^ order branches closest to the cell body, 2^nd^ order branches the ones after the first ramification point, etc). All data were expressed as Mean±SEM and tested for significance by unpaired t-tests or Mann-Whitney tests as appropriate using GraphPad Prism 6.0.

## Acknowledgements

Supported by grants from the BBSRC (AB, Grant Number BB/M029379/1), MRC (AB, Grant Number MR/P025811/1), Alzheimer’s Research UK (DG-N, AB, Grant Number PG2014B-2), University of Portsmouth PhD Programme (AB, ICR), CNR – Joint Laboratories (MC, LR), and Programme Avenir Lyon Saint-Etienne (OR).

## Author Contributions

*Irene Chacon-De-La-Rocha*: Formal Analysis; Investigation; Methodology; Writing - original draft. *Gemma Fryatt*: Formal Analysis; Investigation; Methodology; Validation. *Laura Restani*: Investigation; Methodology. *Andrea D. Rivera*: Investigation. *Matteo Caleo, Olivier Raineteau*: Resources, Writing – review & editing. *Diego Gomez-Nicola:* Conceptualization; Data curation; Formal analysis; Funding acquisition; Project administration; Resources; Supervision; Validation; Visualization; Writing - review & editing. *Arthur M. Butt*: Conceptualization; Data curation; Formal analysis; Funding acquisition; Project administration; Resources; Supervision; Validation; Visualization; Writing - original draft; Writing - review & editing.

## Competing Interests

Prof Arthur Butt and Dr Andrea Rivera are shareholders and co-founders of the company GliaGenesis. All authors declare no other conflicts.

## Data Availability

All data generated or analysed during this study are included in this published article.

## Notes

### Competing Interest Statement

The authors have declared no competing interest.

